# The Emergent Connectome in *Caenorhabditis elegans* Embryogenesis

**DOI:** 10.1101/146035

**Authors:** DevoWorm Group, Bradly Alicea

## Abstract

The relatively new field of connectomics provides us with a unique window into nervous system function. In the model organism *Caenorhabditis elegans*, this promise is even greater due to the relatively small number of cells (302) in its nervous system. While the adult *C. elegans* connectome has been characterized, the emergence of these networks in development has yet to be established. In this paper, we approach this problem using secondary data describing the birth times of terminally-differentiated cells as they appear in the embryo and a connectomics model for nervous system cells in the adult hermaphrodite. By combining these two sources of data, we can better understand patterns that emerge in an incipient connectome. This includes identifying at what point in embryogenesis the cells of a connectome first comes into being, potentially observing some of the earliest neuron-neuron interactions, and making comparisons between the formally-defined connectome and developmental cell lineages. An analysis is also conducted to root terminally-differentiated cells in their developmental cell lineage precursors. This reveals subnetworks with different properties at 300 minutes of embryogenesis. Additional investigations reveal the spatial position of neuronal cells born during pre-hatch development, both within and outside the connectome model, in the context of all developmental cells in the embryo. Overall, these analyses reveal important information about the birth order of specific cells in the connectome, key building blocks of global connectivity, and how these structures correspond to key events in early development.

## INTRODUCTION

Connectomics can tell us a lot about the way behavior is generated from networks of neurons such as the central nervous system or tissues such as the retina (Seung, 2012; Helmstaedter et al, 2013). In mammalian brains, connectomes at cellular resolution are difficult to define and interpret (Morgan and Lichtman, 2014; Sporns et al, 2005). Yet in the nematode *Caenorhabditis elegans*, the connectomics of a 302 cell nervous system can be easily visualized and analyzed. With its relatively small size, single-cell specificity, and easily-observed physical connections, the *C. elegans* connectome serves as a model for function relative to more complex nervous systems (Chatterjee and Sinha, 2007). As such, data describing *C. elegans* connectivity are easily accessible (see Methods).

The *C. elegans* connectome has been formally-defined as a connectivity matrix by Jarrell et al (2012) and Varshney et al (2011). Previous attempts at characterizing the connectome in *C. elegans* has focused on constructing a model of the static adult version based on chemical (synaptic) or physical (cell-cell interactions) relationships. Yet we can combine adult connectome data with cellular data from embryogenesis to yield a dynamic version of the connectome as it emerges from developmental cells. While there are significant anatomical differences between neuronal cells in the embryo and the adult version of the connectome, our analysis is nevertheless informative of how the cells associated with the adult connectome emerges during embryogenesis.

In *C. elegans*, we can also distinguish between developmental cells and terminally-differentiated cells. In this paper, we conduct analysis of both developmental cells and terminally-differentiated cells, with only the latter considered as part of the connectome. Although separate from the predictions of a connectome, we can also create a connectivity map between developmental cells by establishing a distance-based network by on potential spatial relationships such as intercellular signaling or structural differentiation (see Alicea and Gordon, 2018). Analysis of developmental cells such as these can help to provide insights into patterns found in structural and perhaps even functional patterns of the emerging adult phenotype. Developmental cell lineages of *C. elegans* are established by founder cells (Sulston et al, 1983). The first two founder cells are called AB and P1, and represent developmental cell divisions originating in the anterior and posterior halves of the embryo, respectively. Aside from a few exceptions, developmental cells almost always divide into two daughter cells after a period of time (Wasserstrom et al, 2018). By contrast, terminally-differentiated cells have stopped dividing and can be identified by their adult nomeclature and function. Terminally-differentiated cells also have a direct developmental cell ancestral lineage, and because of this we can analyze these cell lineages to detect relationships between early embryogenesis and later development or even adulthood.

By investigating developmental cell lineages and other features of the developmental milieu, we may be able to identify the origin of modules and functional subdivisions of the connectome (Kaiser, 2011; Reigl et al, 2004). By using *C. elegans* as a model organism, we can also make connections between developmental cell lineages and adult neuronal network topologies. One benefit of combining multiple datasets is to investigate previously unexplored areas of inquiry not possible by relying on a single dataset or model (Faisal et al, 2014). Making an explicit link between the emergence of neuronal cells in embryogenesis and the adult connectome provides clues as to which relationships in the embryo (such as the spatial proximity of developmental cells, birth order of terminally-differentiated neuronal cells) also give rise to neuronal cell-neuronal cell connectivity (Varier and Kaiser, 2011).

In this paper, we will investigate the origins of *C. elegans* connectome in the embryo using a complex network approach. Then, we will combine previously published connectome (Varshney et al, 2011) data with a bioinformatic approach (Alicea et al, 2018) to establish the connections between the process of developmental cell differentiation and terminally-differentiated cells emerging into a coherent nervous system. In this way, we contribute to the literature an analysis based on analyzing a previous unexplored combination of data sources (cell annotations, cell lineage tracing, and a connectome based on gap junction connections between neuronal cells). This will yield a series of time-dependent networks in addition to insights regarding how the embryogenetic process influences the formation of functional neuronal networks. This work also complements work by Nicosia et al (2013), in which the connectivity between neuronal cells in the newly-hatched embryo demonstrates an economy of wiring as the worm transitions from embryo to first larval stage.

Our analytical strategy will focus on three phenomena related to the emergence of the connectome in development. The first of these is investigating the temporal emergence of neurons in the embryo. Using data on birth times for terminally-differentiated neurons, we can plot the first and all subsequent neuronal birth times according to which neurons have been born at or before 265, 280, 290, 300, and 400 minutes post-fertilization. Yet this only provides us with lists of cells. Therefore, we investigate the developmental cell lineages from which the neuronal cells derive. This is done by linking the *C. elegans* developmental cell lineage to a model of the adult connectome and neuronal birth time data in general. Aside from the potential to construct embryo networks from patterns the emerge from differentiation and geometric organization (Alicea and Gordon, 2018), such a linkage also demonstrates the potential role of developmental sublineage distribution on subsequent connectome organization. Finally, the phenotypic context of the proposed embryogenetic connectome is examined. By placing the neurons born during embryogenesis in the spatial context of developmental cell lineages, we can begin to appreciate the spatial transformations that must occur as the neurons go from existing in a spherical mass to becoming part of a fully-functioning nervous system.

## METHODS

### Developmental Cell Lineage Data

Datasets consist of pre-hatching (up to 558 cells) cell divisions (Bao et al, 2008). During acquisition of these data, cell nucleus identification was done by identifying GFP+ markers using the Starry Night software package. Three-dimensional (spatial) position and nomenclature (identity) for all cells are extracted in this manner from 261 embryos. The position of each cell is averaged across all embryos along each anatomical axis (anterior-posterior, left-right, and dorsal-ventral). The positional data is then translated to have its origin at the location of the P0 cell, which produces a three-dimensional spatial representation of the embryo.

### Embryonic Time Series

To construct the embryonic time series of terminally-differentiated neurons, timed cell lineage data were acquired courtesy of Nikhil Bhatla and his Interactive lineage tree application (http://wormweb.org/celllineage). More information regarding our use of these data to build a embryonic time-series can be found in Alicea et al (2018). Cells represented in an embryo at a given time are determined by first calculating the lifespan of each cell in the lineage tree (e.g. the time at which each cell is born and each cell either divides or dies), and then identifying all cells alive at a given time. Terminally-differentiated cells were assumed never to die, unless specified by the data. These data were also used to establish a link between developmental cell sublineages and terminally-differentiated cells.

### Connectome data

The connectome data was provided by the Open Connectome database (http://openconnecto.me/). The connectome data originated with nervous system cells (n = 279) from the adult hermaphrodite (Varshney et al, 2011). The connectivity matrix with electrical synaptic weights were matched against our list of terminally-differentiated cells at 280, 290, 300, and 400 minutes. The resulting cells were used to create unique connectivity matrices for each timepoint, and included unconnected cells. These data were visualized in Gephi 0.9.0 and modeled using yED. Adjacency matrices were also created from the connectivity information. All files (r code, GraphML code, and data files) are available at https://github.com/devoworm/embryogenetic-connectome.

The adjacency matrix data provided in Varshney et al (2011) was extracted from previously collected electron micrographs (White et al, 1986). The network adjacency matrices in this paper are defined and weighted by electrical synapses called gap junctions (Majewska and Yuste, 2001), but is ordered by the additional existence of chemical synapses. The gap junction-based weights are determined by the number of gap junctions shared between a pair of cells (Varshney et al, 2011).

### Developmental connectome modeling in adult phenotype

The lateral-view digital model of the hermaphrodite adult *C. elegans* using the Supplemental Figure 1 and Supplemental Figure 2 originates from the Virtual Worm project. The original model was constructed using Blender (available at http://canopus.caltech.edu/virtualworm/Virtual%20Worm%20Blend%20File/), and the annotated Blender model (with connectome data included as a series of scenes) is available at https://github.com/devoworm/embryogenetic-connectome/blob/master/developmental-connectome-265-400-minutes-adult-model.zip

**Figure 1.**
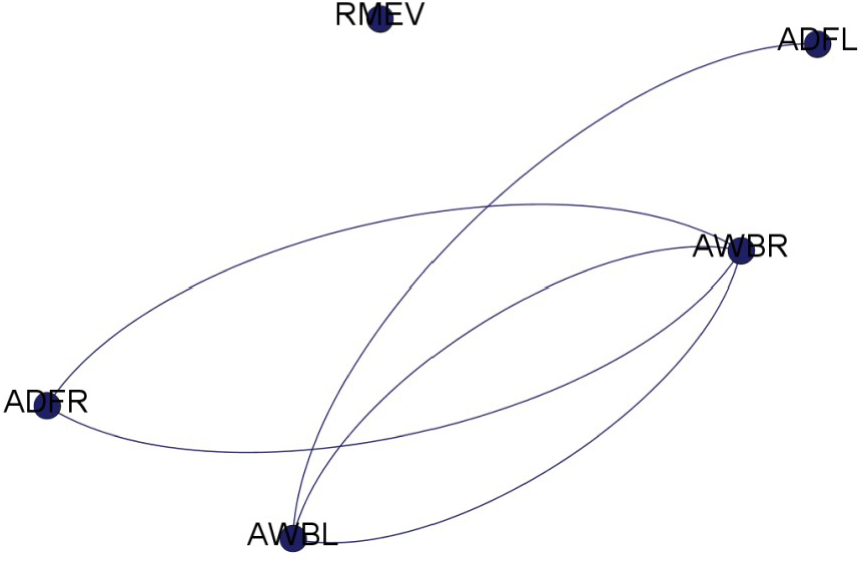
*C. elegans* embryonic connectome at 280 minutes. Blue nodes are all terminally-differentiated cells born by 280 minutes of embryogenesis.

**Figure 2.**
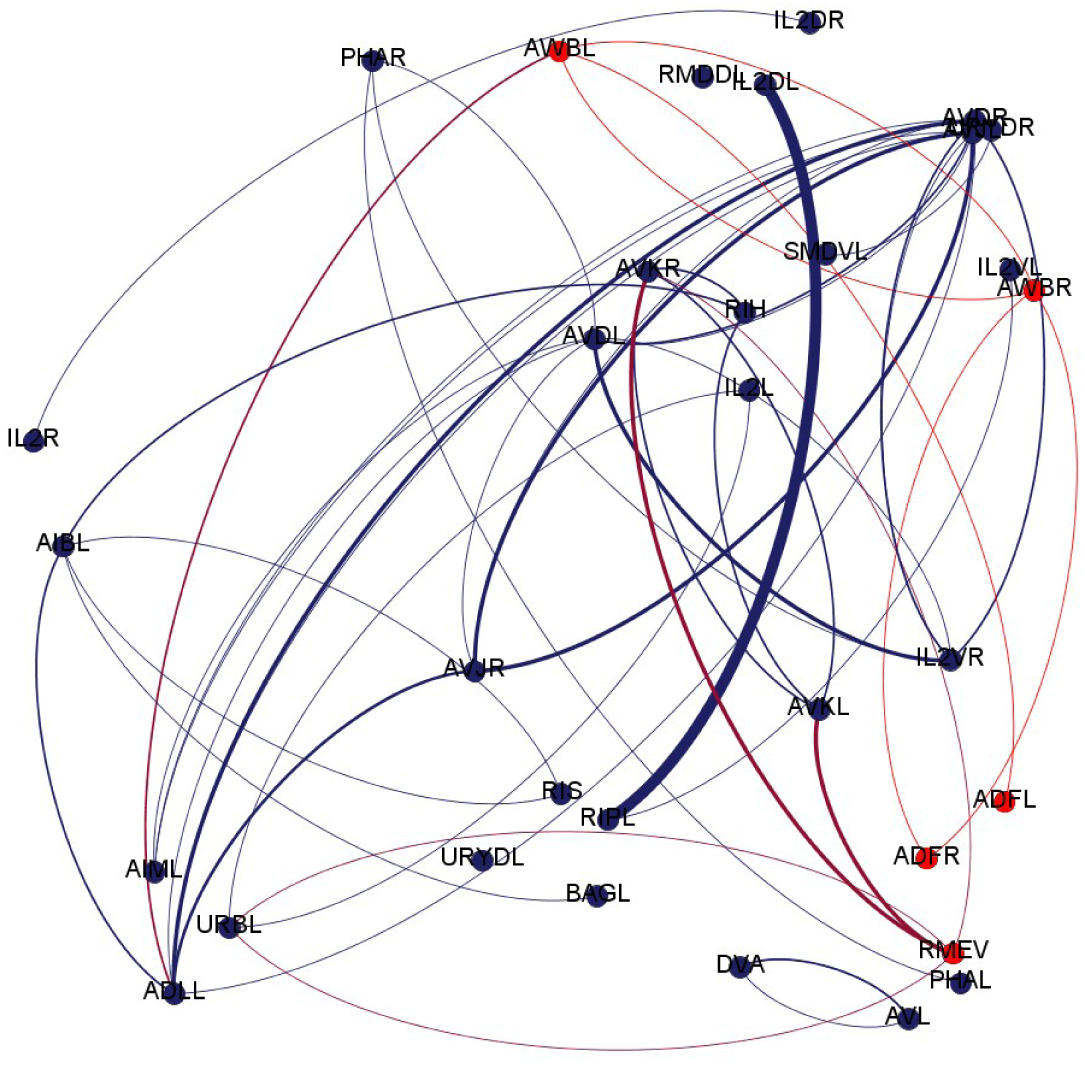
*C. elegans* embryonic connectome at 290 minutes. Red nodes are all terminally-differentiated cells born by 280 minutes of embryogenesis, and blue nodes are all terminally-differentiated cells born by 290 minutes of embryogenesis.

## RESULTS

### Temporal Emergence of Neurons in the Embryo

Once developmental cells from the AB lineage terminally differentiate, some of them become neurons. By combining connectomic data from Varshney (2011) and embryo birth time time-series from Alicea et al (2018), we construct connectome for three time points of interest: 280, 290, and 300 minutes of embryogenesis. These time points were selected for their emphasis on the emergence of neuronal cells and their connectivity during this time period. As has just been demonstrated, only developmental cells are generated before 200 minutes. Yet by 400 minutes, all terminally differentiated cells not generated post-embryonically are also present in the developmental time-series.

Figure 1 demonstrates the first stage of connectivity in the connectome. The earliest coherent connectome (cells with synaptic connectivity) emerges at 280 minutes of embryogenesis. The first neuronal cell to emerge in the connectome (RMEV at 265 minutes) is also disconnected from a network of five other neurons at 280 minutes. These four connected neurons reveal another interesting attribute of early connectomes. Neuronal cells often emerge in pairs (e.g. symmetrical components), and become reciprocally interconnected. In the vast majority of cases, these involve left-right pairs. Bilateral reciprocity becomes a motif in these early networks, and a means to build more complex connectivity patterns later on.

Figure 2 and 3 show the embryonic connectome connectome at 290 and 300 minutes, respectively. Aside from denser and more complex connectivity patterns, the weights of synaptic connection increase while also exhibiting greater variation during and after 300 minutes. One-way and reciprocal binary connectivity now combine to form motifs while also serving as a means to build more complex connectivity patterns later on. For example, while RMEV is unconnected at 280 minutes, RMEV becomes connected to several neurons emerging at 290 minutes. This increase in connectivity is enough for RMEV to become a regional hub in the network.

**Figure 3.**
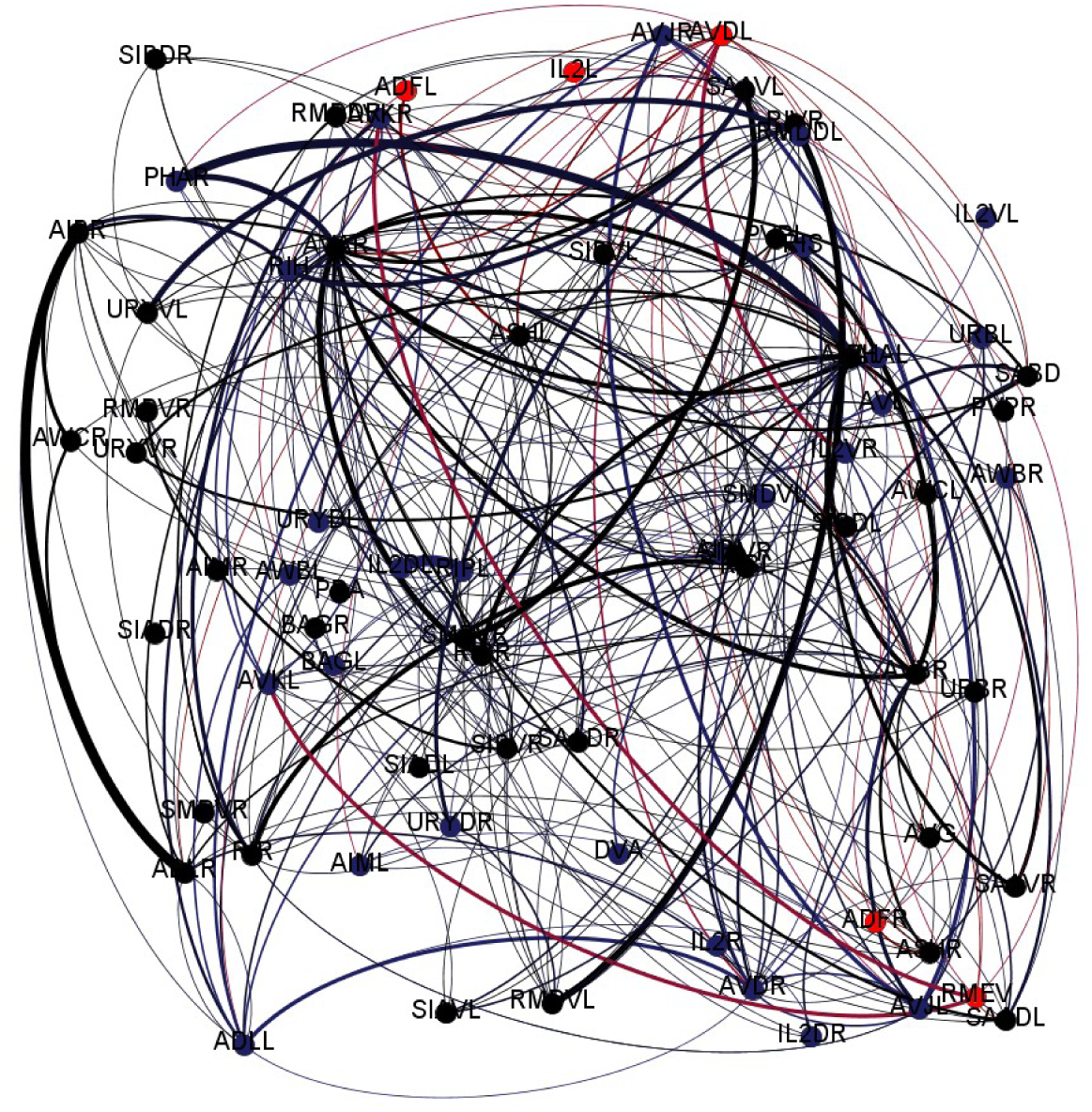
*C. elegans* embryonic connectome at 300 minutes. Red nodes are all terminally-differentiated cells born by 280 minutes of embryogenesis, black nodes are all terminally-differentiated cells born by 290 minutes of embryogenesis, and blue nodes are all terminally-differentiated cells born by 300 minutes of embryogenesis.

To better understand how motifs and patterns of connectivity are formed in the embryonic connectome over time, several proportions are calculated for four timepoints (280, 290, 300, and 400 minutes). The results are shown in Table 1, and looking at the trends for reciprocal connections per timepoint and proportion of source and target nodes per timepoint yields an interesting relationship. At 280 minutes, we observe a few reciprocally-connected neurons that make up the entirety of the network. By 290 minutes, about 30 additional neurons are born, many of which only have an input or output, and many of those lacking a bilateral counterpart. At 300 minutes, many these cells gain input and/or output connections that make them more fully functioning members of the network. Once we reach 400 minutes, many more cells are born that serve as bilateral counterparts to existing cells. Thus, the 280-300 minute period is defined by asymmetric connectivity patterns, while from 300 minutes on network connectivity becomes more complex (Azulay et al, 2016), as feedback loops and large-scale network components are assembled.

**Table 1.** Number of reciprocal connections (ordered connections from cell A to cell B and from cell B to cell A) and nodes connected to network serving as both sources and targets.

|  | **280** | **290** | **300** | **400** |
| --- | --- | --- | --- | --- |
| **Number of nodes** | 4 | 31 | 69 | 163 |
| **Reciprocal Connections** | 0.8 | 0.46 | 0.39 | 0.74 |
| **Source and Target Nodes** | 0.75 | 0.61 | 0.9 | 0.98 |

### Neuronal Origins in Developmental Cell Lineages

The embryonic connectome can be analyzed in two ways: a dynamic version, and a static version that explicitly demonstrates linkages with developmental cell lineages. To render the latter, we took all cells in the connectome dataset and matched them in two ways. The first match involved taking all cells in the connectome dataset and matching them to cells from the 300 minute point in our embryogenetic time series. The second match involved taking each cell resulting from this intersection operation and linking to its corresponding 8-cell embryo ancestor representing the developmental cell lineage. This resulted in a classification of cell matches by one-way versus reciprocal connectivity. A quick summary of these results are shown in Table 2.

**Table 2.**
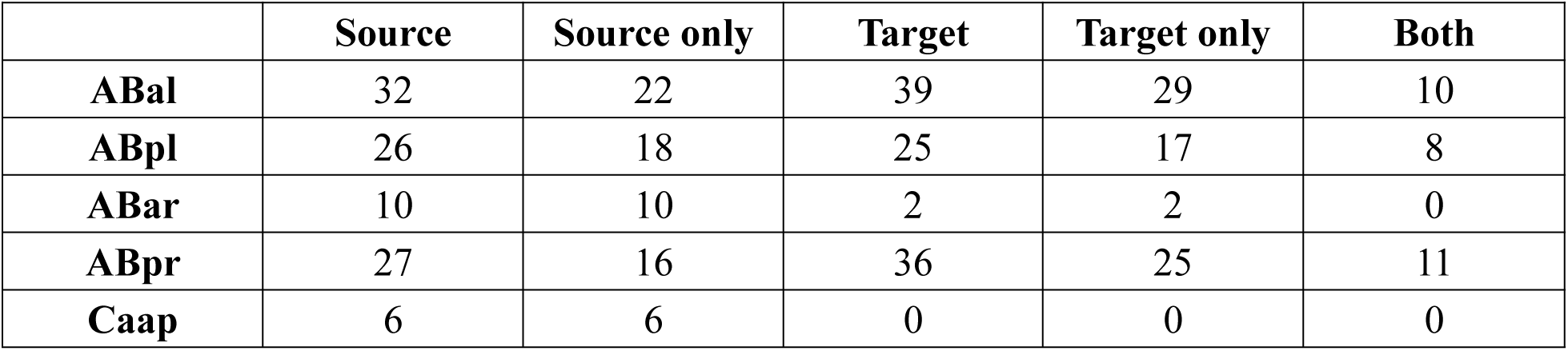
Characterization of connections amongst neurons by developmental subtree of origin emerging between 300 and 400 minutes of embryogenesis. Network generated using a synaptic <u>threshold of 4.0.</u>

|  | Source | Source only | Target | Target only | Both |
| --- | --- | --- | --- | --- | --- |
| **ABal** | 32 | 22 | 39 | 29 | 10 |
| **ABpl** | 26 | 18 | 25 | 17 | 8 |
| **ABar** | 10 | 10 | 2 | 2 | 0 |
| **ABpr** | 27 | 16 | 36 | 25 | 11 |
| **Caap** | 6 | 6 | 0 | 0 | 0 |

A visual map of these the results in Figures 4 and 5. Figure 4 shows the connectome for four subtrees at 300 minutes of embryogenesis. In Figure 4 (left) we can see the full connectivity of this connectome color-coded by subtree, while in Figure 4 (right) only connections above a nominal threshold are shown. In the thresholded case (Figure 4, right), some cells retain their connectivity and serve as network hubs, while other node become disconnected. Figure 5 shows the entire connectome network stratified into four subnetworks based developmental cell sublinege (ABal, ABpl, ABar, and ABpr).

**Figure 4.**
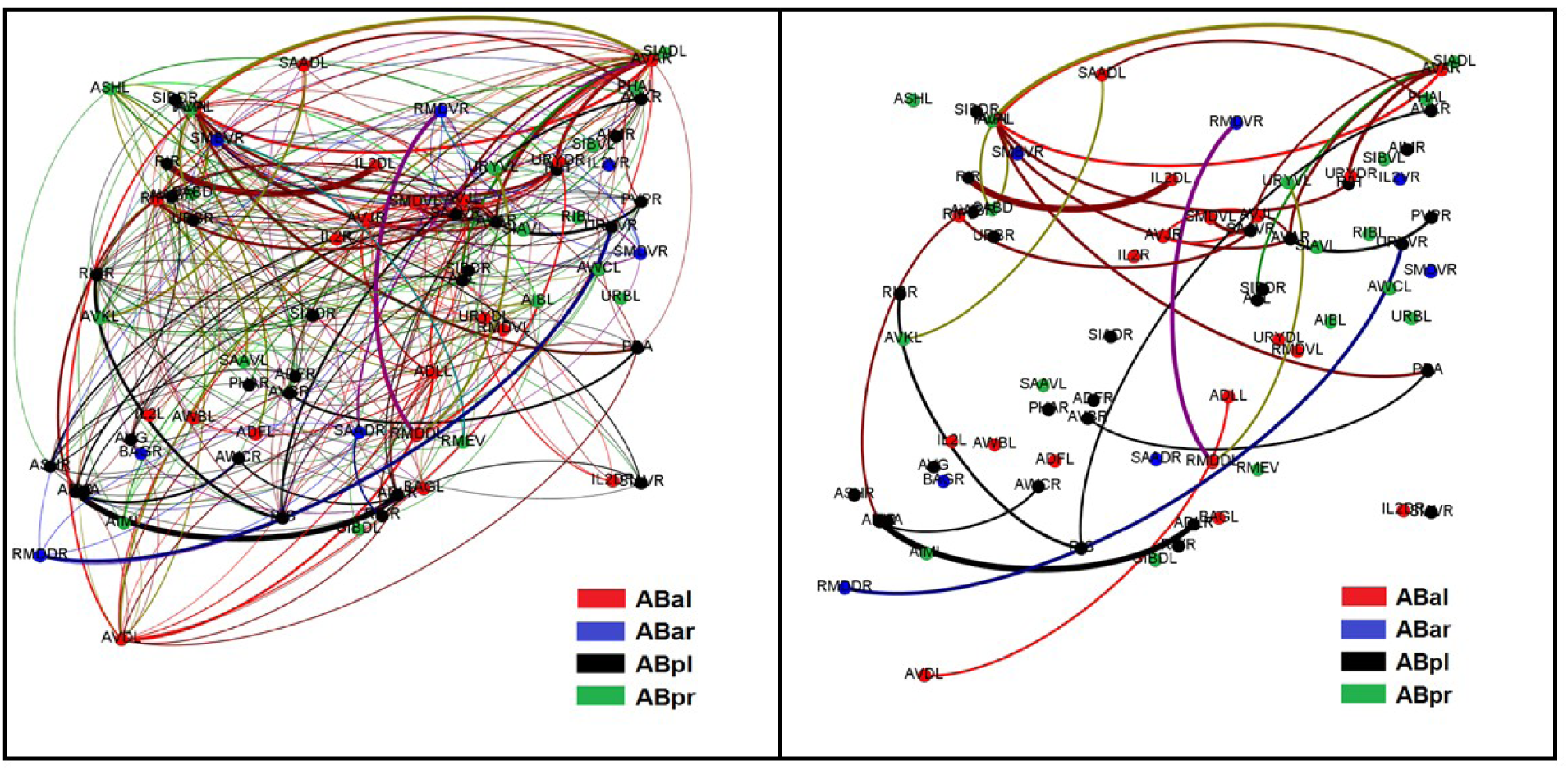
*C. elegans* embryonic connectome at 300 minutes of embryogenesis. LEFT: entire connectome, nodes are color coded by developmental cell sublineage of origin. RIGHT: edges with a value greater than 4.0, nodes are color coded by developmental cell sublineage of origin.

**Figure 5.**
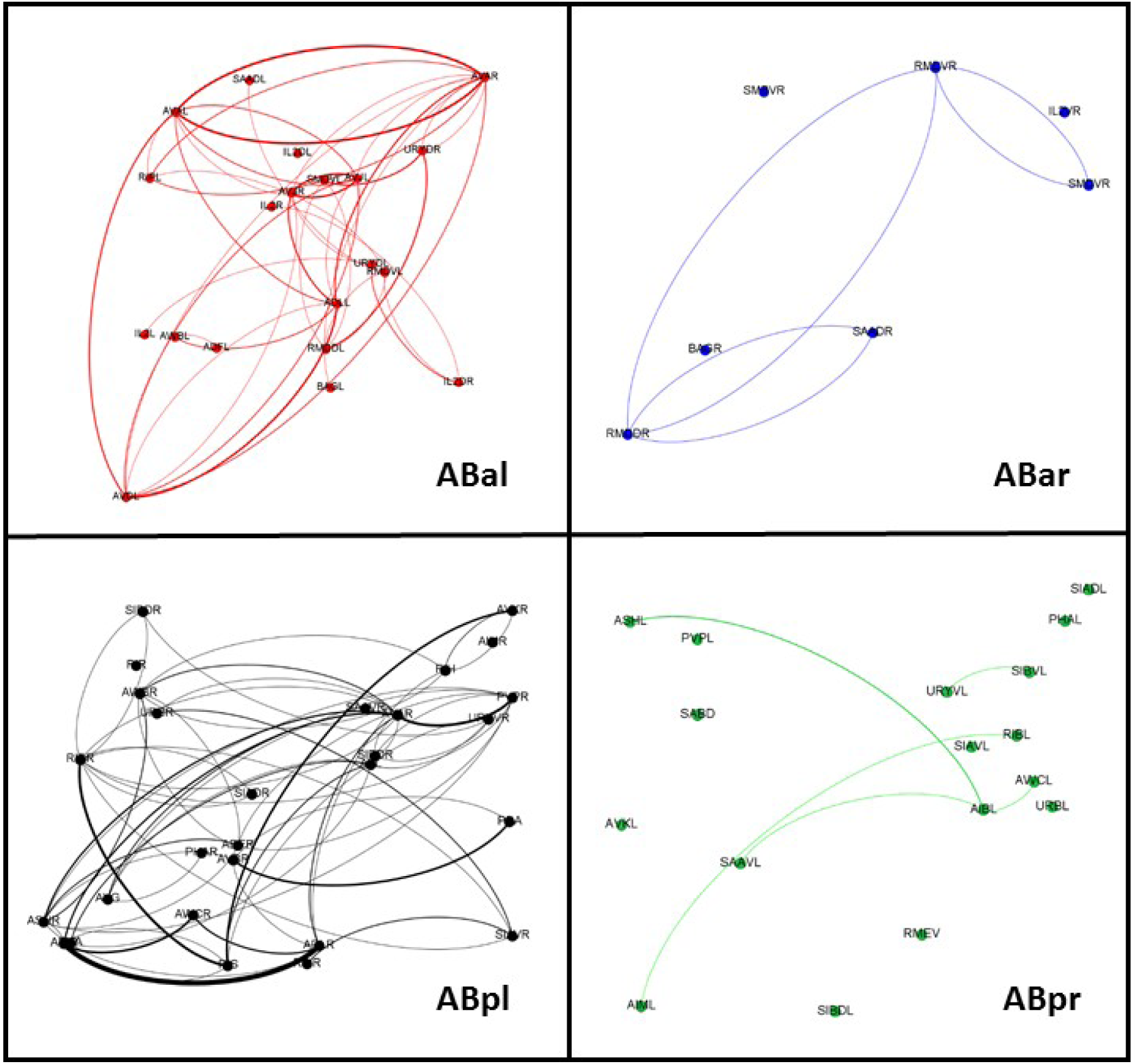
*C. elegans* embryonic connectome at 300 minutes of embryogenesis by developmental cell of origin. Terminally-differentiated cells that descend from four cells in the 8-cell embryo.

To better understand the emerging function and topology of the *C. elegans* connectome, data from this analysis was compared to a list of so-called “rich club” neurons (Towlson et al, 2013). The results shown in Table 3 and demonstrate the relationship (or lack thereof) between adult position in the connectome and embryogenetic origins. Despite all sharing the rare attribute of being (connection) rich in adulthood, there appears to be no single developmental recipe to their success. The list of developmental cell lineage origins in Table 3 demonstrates almost an even representation of cells originating in the anterior and posterior components of the AB sublineage. Additionally, while rich-club neurons come from a range of birth times, they are all born at or before 350 minutes of embryogenesis.

**Table 3.** Birth times, developmental lineage of origin, and identity by annotation for 14 ‘rich club’ terminally-differentiated neuronal cells as defined in Towlson et al (2013).

| Cell Name | Neuron ID | Birth Time (min) | Developmental Lineage | Annotation |
| --- | --- | --- | --- | --- |
| AVAR | AS9 (P9.apa) | 295 | ABal | Ventral Cord Motorneuron |
| AVAL | AS8 (P8.apa) | 295 | ABal | Ventral Cord Motorneuron |
| AVBR | ASER | 300 | ABpr | Ventral Cord Interneuron |
| AVBL | ASEL | 350 | ABpl | Ventral Cord Interneuron |
| AVDR | ASGR | 290 | ABal | Ventral Cord Interneuron |
| AVDL | ASGL | 290 | ABal | Ventral Cord Interneuron |
| AVER | ASHR | 350 | ABpr | Ventral Cord Interneuron |
| AVEL | ASHL | 350 | ABal | Ventral Cord Interneuron |
| PVCR | --- | 350 | ABpr | Ventral Cord Interneuron |
| PVCL | --- | 350 | ABpl | Ventral Cord Interneuron |
| DVA | CP9 (P11.aapp) | 280 | ABpr | Ring Interneuron |
| AIBR | AIAR | 295 | ABpr | Amphid Neuron* |
| RIAR | --- | 300 | ABal | Ring Interneuron* |
| RIBL | --- | 295 | ABpl | Ring Interneuron* |

\* neurons in the 1 $\sigma$ and 2 $\sigma$ group, but not in the main 11.

Given the concurrence of neuronal and developmental information, there are a few additional observations. First, most connectome cells at 300 minutes of embryogenesis derive from either the ABal or ABpl lineage. Secondly, neurons that derive from the ABal developmental sublineage begin to form a clique with a core of connected cells. To understand the role of developmental cell lineage of origin in connectivity at 300 minutes, we can look at descendants of each developmental cell sublineage to look at patterns of connectivity (Figure 5).

Using the full set of connectome information, we can see in Figure 5 that disconnected cells exist among descendants of sublineages ABar and ABpr only. The interconnections amongst these subnetworks are less dense than subnetworks representing ABal and ABpl. When the connectome is thresholded, cells from all sublineages (ABal, ABar, ABpl, and ABpr) become disconnected (Figure 4, right). However, a terminally differentiated cell’s time of birth has no bearing on connectivity patterns.

### Phenotypic Context of the Embryogenetic Connectome

We also visualize the data to gain more information about the locations of these cells in context of the adult phenotype. We can model the connectome using a lateral-view model of the adult hermaphrodite across four developmental sublineages of origin (ABal, ABar, ABpl, ABpr) at 300 minutes (Supplemental File 1) and over embryogenetic time from 265 to 300 minutes post-fertilization (Supplemental File 2). While the lateral view shows the relative positions of cells along an orientation representing the lineage tree, this model also obscures spatial structure along the left-right axis.

To gain a better understanding of the link between the unfolding of spatial organization in the embryo and potential structural biases present in adult phenotypes, we used the data shown SF1 and counted all cells that are part of a bilateral pair. We then assessed their positional laterality as terminally-differentiated cells. For the ABar subtree, all cells (7 of 7) are on the right-hand side. The ABpl subtree yields 20 of 28 cells on the right-hand side and 1 of 18 cells on the left-hand side. The remaining AB sublineages are biased towards the left-hand side: in ABpr, 15 of 17 cells are on the left-hand side, while ABal contains 15 of 20 cells on the left-hand side and only 5 of 20 cells on the right-hand side. While there does seem to be a relationship between subtree origin and structural modularity, we cannot make any more significant statements about these findings.

One issue with using a connectome observed in adulthood to infer the embryogenetic emergence of connectivity is lack of information about spatial context in the embryo itself. To gain insight into transformations between the embryo and adult phenotypes, we traced the terminally-differentiated neurons back to the embryonic locations of their developmental precursor cells. This provided positional information for the neuronal cells, and allows us to embed these cells within a three-dimensional visualization of the embryo. Supplemental File 3 provides a time series of neuronal cells. Each featured time point presents three sets of images. The first set of images shows the neuronal cells present in the embryo at a specific sampling point for the following times post-fertilization: 265, 280, 290, 300, and 400 minutes. The second set of images shows each of these neurons embedded in a cell centroid model of the embryo, based on the location of their immediate developmental cell precursor. Finally, the transition between each sampling point shows all cells born during this interstitial period.

Figure 6 shows us what the relationship between neuronal and developmental cells looks like from multiple perspectives. Figure 6a is a three-dimensional plot of all developmental cells and neuronal cells in the embryo regardless of birth time. This reveals a result similar to Nicosia (2013) in that two distinct spatial patterns for emerging neurons: a population of neurons near the floor of the embryo, and a population of neurons scattered about the midsection of the embryo. Table 4 shows the distribution of cell types in each population based on functional annotations of cells in each population. According to these criteria, spatial subdivision does not seem to be related to differences in adult function, which might reflect the pre-establishment of localized anatomical structures. Figure 6b is a two-dimensional plot (excluding the dorsal-ventral axis) that shows a top-down view of neurons embedded in the embryo. This reveals another spatial pattern: the floor and the midsection populations are oriented orthogonally to one another. Figure 6c pulls this together in the context of the adult (independent of birth time), and reveals that the midsection neurons go on to become neurons in the adult head, while floor neurons go on to be neurons in the adult midsection and tail.

**Table 4.**
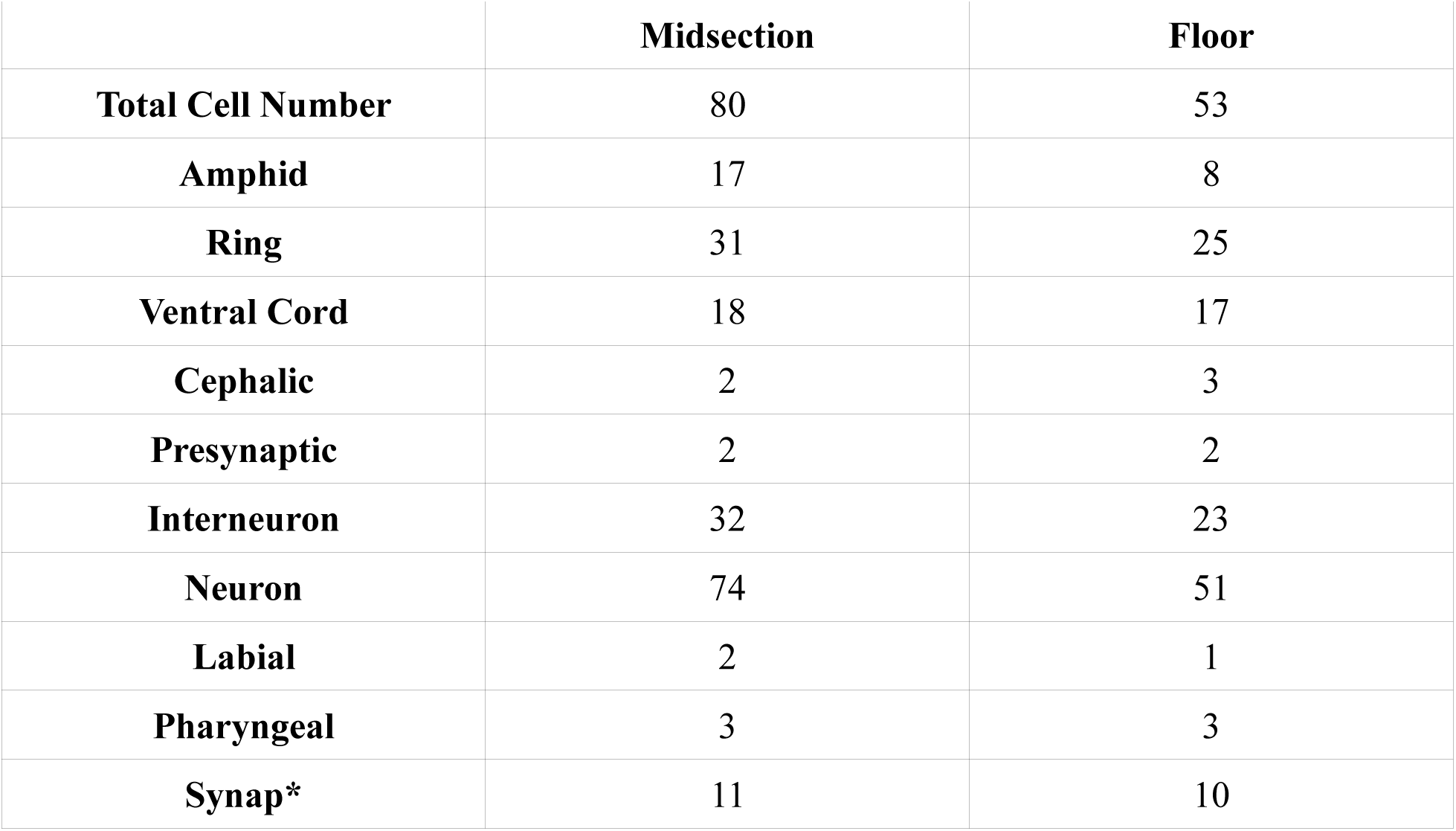
Total number of cells in the midsection and floor populations of neurons present in the embryo, and occurrence of ten most common terms in the list of annotations for all cells in each population. Wildcard (*) represents all instances of a root term. Total cell number does not equal total number of occurrences of all terms for each population.

**Figure 6.**
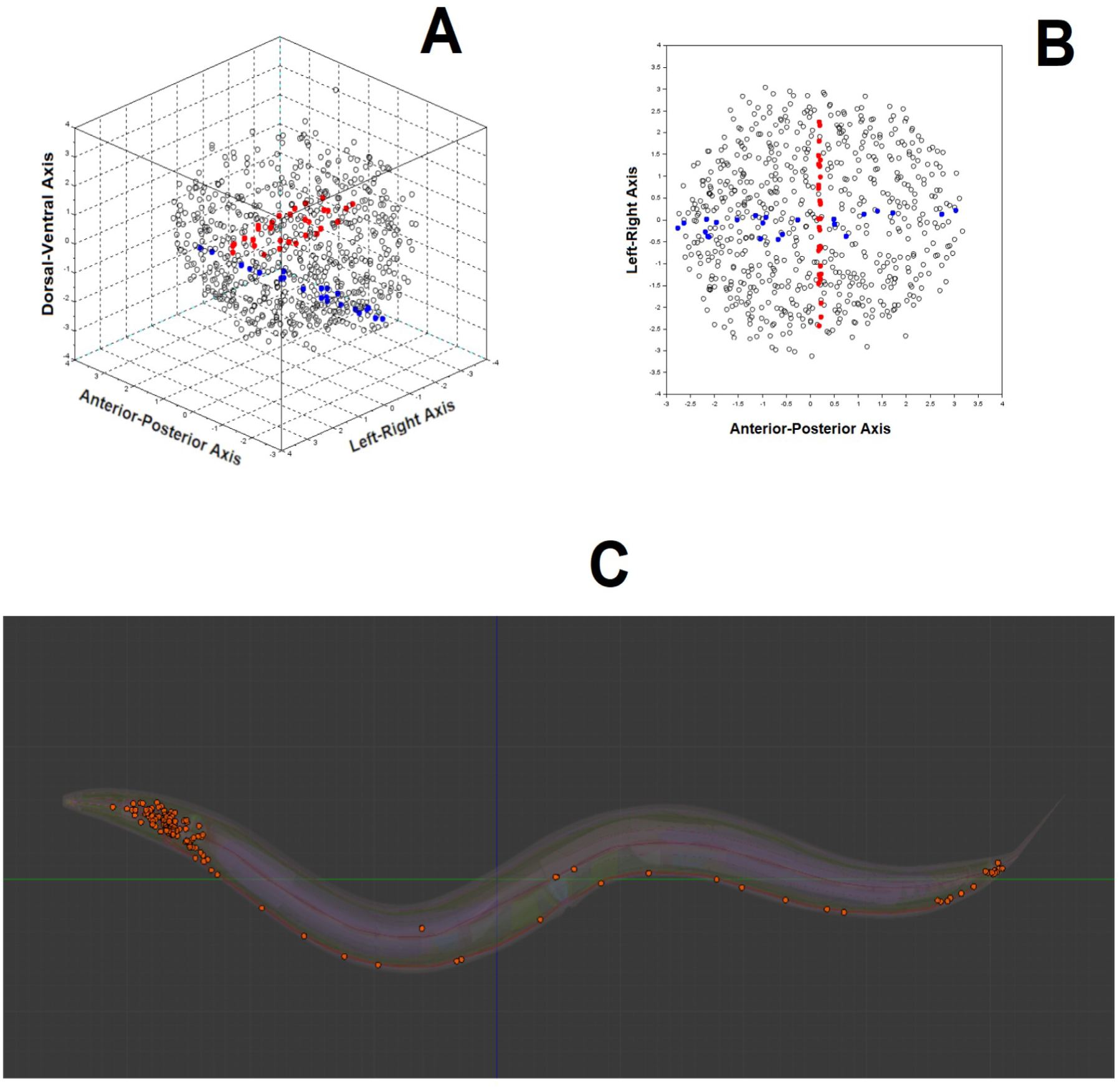
Three ways of visualizing all neurons born during embryogenesis (pre-hatch development). Neuronal position is based on the x,y,z location of their immediate ancestor. This map is model-free with respect to a formal connectome. A: neurons and developmental cells in a three-dimensional representation of the embryo. B: neurons and developmental cells in a two-dimensional (top-down) view of the embryo, anterior-posterior axis (x) versus left-right axis (y). In both A and B, developmental cells are represented by black circles, floor population of neurons are filled with blue, midsection neurons are filled with red. C: neurons (orange) plotted in A and B with their position in the adult hermaphrodite.

## DISCUSSION

We have been able to demonstrate the developmental origins of the connectome, a network that emerges from the first terminally-differentiation neurons in *C. elegans* and is shaped by both their ancestral developmental cell lineages and proximal relationships between these cells. There is also a relationship between network topologies representing developmental and terminally-differentiated cells. This involves a comparison between developmental cell lineages and the emergence of terminally-differentiated cells. Since linkages in this particular connectome model is determined by the existence of gap junctions between cells, there should be some degree of similarity what is inferred for the embryo and the adult connectome.

Our connectome data are bounded by the 200 to 400 minute interval, which is the time between the first appearance of non-germline terminally differentiated cells and the comma stage of development (Chisholm and Hardin, 2005). The embryonic connectome first emerges at 280 minutes, and grows quickly up to the 400 minute mark. Within this span of time, the embryonic connectome transitions from a simple interconnected set of neurons to a complex network. In the original analysis of the connectome data (Varshney et al, 2011), up to 11 types of motifs were identified in the full connectome based on gap junctions. While we did not conduct a motif analysis on these data, we can still observe rudimentary motifs and related patterns, particularly the emergence of reciprocally connected pairs, disconnected cells, and even preferential attachment (Betzel et al, 2016).

One way to interpret these results is to consider what is happening at 280 to 400 minutes of embryogenesis, and in particular the interval from 280 to 300. We have observed that the first proto-connectome emerges at 280 minutes of embryogenesis. We have also observed that the emergence of hub neurons occurs between 290 and 300 minutes of embryogenesis. During this brief period of time, the embryo undergoes its first cleavage at 280 minutes (Soto et al, 2002) with dorsal intercalation and ventral cleft closure occurring by 290 minutes (Sulston et al, 1983; Altun and Hall, 2011). The conventional wisdom is that all connectome cells are generated during the first proliferation phase (Altun and Hall, 2011). We can refine this finding by suggesting that the connectome first proliferates between 280 and 300 minutes, and continues to grow in size until the 400 minute mark. At this point in time, the connectome appears to be structurally complete. This is consistent with the morphogenesis of the pharynx, which forms on a similar timescale according to the rules of multi-step morphogenesis (Portereiko and Mango, 2001). Other parts of the adult phenotype also begin to exhibit their adult form during this period. For example, at the 310 minute mark, cell migrations such as the movement of dorsal hypodermal nuclei to the side of the midline opposite from their origin are taking place (Chisholm and Hardin, 2005). The 400 minute mark is also the beginning of the elongation phase and the end of the comma phase (Chisholm and Hardin, 2005).

There are a few caveats to consider, particularly in terms of pursuing future directions. One issue is that neuronal birth time is not particularly informative of the details of later synaptic connectivity (Kratsios et al, 2015). It has previously been found that early-born neurons tend to be both more highly connected and exhibit more longer-range connections than later-born neurons (Varier and Kaiser, 2011). Yet neuronal birth time along with their developmental cell ancestry may be informative of later aggregate patterns in the *C. elegans* nervous system. As most neurons are present before hatching, contacts between adjacent neurons in early development might be important in establishing long-distance connectivity patterns (Varier and Kaiser, 2011). Future work might take into account the role of axonal projections and cell migrations in the process of assembling the adult connectome.

## ACKNOWLEDGEMENTS

We would like to acknowledge all primary data donors (datasets specified in the Methods section), without which this paper would not be possible. Thanks also go to the OpenWorm Foundation, which provided institutional and intellectual support, as well as Dr. Oliver Hobart, who offered some useful advice. Additional thanks go to past and current members of the DevoWorm group, information about which can be found at http://devoworm.weebly.com.

## SUPPLEMENTAL MATERIALS

Supplemental File (Figure) 1. Images of *C. elegans* embryonic connectome cells (shown in blue) born by 300 minutes embryogenesis in the adult hermaphrodite *C. elegans* phenotype. From top: ABal, ABar, ABpl, and ABar sublineages.

Supplemental File (Figure) 2. Images of *C. elegans* embryonic connectome (shown in orange) cells born at the given time in embryogenesis in the adult hermaphrodite *C. elegans* phenotype. From top: 265, 280, 290, and 300 minutes.

Supplemental File (Movie) 3. Animation showing expansion of the C. elegans embryonic connectome in the context of adult hermaphrodite C. elegans phenotype, nuclear positions of cells in the embryo, and the predicted connectivity amongst cells from 265 to 400 minutes in the expanding connectome. The last still shot shows all pre-hatch neurons (so-called model-free approach which does not rely on adult connectome definitions).

